## Supplementary Information for "Specificity and mechanism of the double-stranded RNA-specific J2 monoclonal antibody"

**Supplementary Tables 1-2**
**Supplementary Figures 1-10**

| Ligand (J2 IgG) | Analyte (dsRNA) | $k_{on}$ (×$10^5$ $M^{-1}s^{-1}$) | $k_{off}$ (×$10^{-3}$ $s^{-1}$) | $K_d$ (nM, kinetics) | $K_d$ (nM, steady state) |
|---|---|---|---|---|---|
| WT | 6-bp | N.D. | N.D. | N.D. | N.D. |
| WT | 10-bp | N.D. | N.D. | N.D. | N.D. |
| WT | 14-bp | 2.3 ± 1.9 | 220.3 ± 27.8 | 1400 ± 800 | 2100 ± 600 |
| WT | 20-bp | 5.9 ± 2.8 | 72.1 ± 2.5 | 150 ± 80 | 120 ± 10 |
| WT | 30-bp | 3.5 ± 5.5 | 10.3 ± 15.3 | 39 ± 10 | 12 ± 28 |
| WT | 40-bp | 5.3 ± 0.2 | 17.2 ± 2.8 | 33 ± 6 | 80 ± 15 |
| WT | 50-bp | N.D. | N.D. | N.D. | 22 ± 8 |
| HC A30P | 20-bp | 2.1 ± 1.9 | 55.1 ± 7. 0 | 28 ± 14 | 120 ± 20 |
| HC Y50A | 20-bp | N.D. | N.D. | N.D. | N.D. |
| HC Y50F | 20-bp | 15. ± 5.3 | 124.7 ± 29.1 | 94 ± 49 | 430 ± 40 |
| HC Y52A | 20-bp | N.D. | N.D. | N.D. | N.D. |
| HC Y52F | 20-bp | 5.8 ± 0.3 | 122.7 ± 7.1 | 210 ± 20 | 220 ± 0 |
| HC Y54A | 20-bp | 5.1 ± 0.2 | 138.0 ± 7.8 | 270 ± 10 | 300 ± 30 |
| HC Y54F | 20-bp | 5.7 ± 0.8 | 140.3 ± 6.8 | 250 ± 40 | 710 ± 170 |
| HC N55A | 20-bp | N.D. | N.D. | N.D. | N.D. |
| HC N101A | 20-bp | N.D. | N.D. | N.D. | N.D. |
| HC N101R | 20-bp | N.D. | N.D. | N.D. | N.D. |
| HC P102A | 20-bp | N.D. | N.D. | N.D. | N.D. |
| HC W104A | 20-bp | N.D. | N.D. | N.D. | N.D. |
| LC Y31A | 20-bp | 8.0 ± 3.3 | 136.8 ± 91.4 | 207 ± 21 | 940 ± 560 |
| LC Y31F | 20-bp | 7.6 ± 1.0 | 97.0 ± 3.5 | 130 ± 20 | 360 ± 60 |
| LC S33R | 20-bp | N.D. | N.D. | N.D. | N.D. |
| LC Y38F | 20-bp | 9.5 ± 0.1 | 79.0 ± 13.4 | 83 ± 14 | 150 ± 20 |
| LC W56A | 20-bp | N.D. | N.D. | N.D. | N.D. |
| LC Y101A | 20-bp | N.D. | N.D. | N.D. | N.D. |
| WT | GC_0%, 30 bp | 5.2 ± 1.9 | 146.7 ± 4.0 | 320 ± 140 | 290 ± 60 |
| WT | GC_3%, 30 bp | 1.2 ± 1.7 | 19.1 ± 24.8 | 190 ± 50 | 220 ± 90 |
| WT | GC_17%, 30 bp | 2.4 ± 0.2 | 16.4 ± 0.8 | 470 ± 30 | 36 ± 22 |
| WT | GC_33%, 30 bp | 1.4 ± 0.2 | 148.3 ± 6.7 | 1100 ± 200 | 460 ± 90 |
| WT | GC_47%, 30 bp | 1.6 ± 1.3 | 19.5 ± 2.9 | 560 ± 810 | 110 ± 20 |
| WT | GC_63%, 30 bp | N.D. | N.D. | N.D. | N.D. |
| WT | GC_77%, 30 bp | N.D. | N.D. | N.D. | N.D. |
| WT | GC_90%, 30 bp | N.D. | N.D. | N.D. | N.D. |
| WT | GC_100%, 30 bp | N.D. | N.D. | N.D. | N.D. |
| WT | Poly (rAU+rAU), 40 bp | 9.2 ± 10.6 | 103.2 ± 30.8 | 310 ± 340 | 91 ± 11 |

**Supplementary Table 1. Summary of BLI analysis parameters.** HC: heavy chain; LC: light chain. N.D.: not determined. $K_d$ (kinetics) was calculated from $k_{on}/k_{off}$ determined by kinetic analyses; $K_d$ (steady state) was derived by curve-fitting sensor responses (nm) as functions of dsRNA concentrations. See Methods for details. Values are mean ± s.d. of three biologically independent samples. All J2 used in BLI analyses are in the form of IgG.

| | J2 Fab bound to 23-bp dsRNA |
|---|---|
| **Data collection** | |
| Space group | $P\ 2\ 2_1\ 2_1$ |
| Cell dimensions | |
| *a*, *b*, *c* (Å) | 65.391 92.374 194.663 |
| α, β, γ (°) | 90, 90, 90 |
| Resolution (Å) | 54.28 – 2.85 (2.90 – 2.85) |
| $R_{sym}$ or $R_{merge}$ | 0.452 (2.77) |
| $R_{pim}$ | 0.085 (0.54) |
| $I / \sigma I$ | 7.3 (1.0) |
| $CC_{1/2}$ | 0.992 (0.444) |
| Completeness (%) | 100.0 (98.1) |
| Redundancy | 29.2 (26.6) |
| **Refinement** | |
| Resolution (Å) | 20.64 – 2.85 (2.96 – 2.85) |
| No. reflections | 28092 (3013) |
| $R_{work}$ / $R_{free}$ | 0.212/0.262 (0.297/0.369) |
| No. atoms | 7309 |
| Macromolecule | 7234 |
| Water | 26 |
| Ligand | 49 |
| *B*-factors | 50.82 |
| Macromolecule | 50.84 |
| Water | 36.61 |
| Ligands | 55.69 |
| R.m.s. deviations | |
| Bond lengths (Å) | 0.002 |
| Bond angles (°) | 0.52 |
| Maximum-likelihood coordinate precision (Å) | 0.36 |
| Protein Data Bank (PDB) accession code | |

**Supplementary Table 2. Summary of data collection and refinement statistics for the co-crystal structures of J2 Fab bound to a 23-bp dsRNA.**

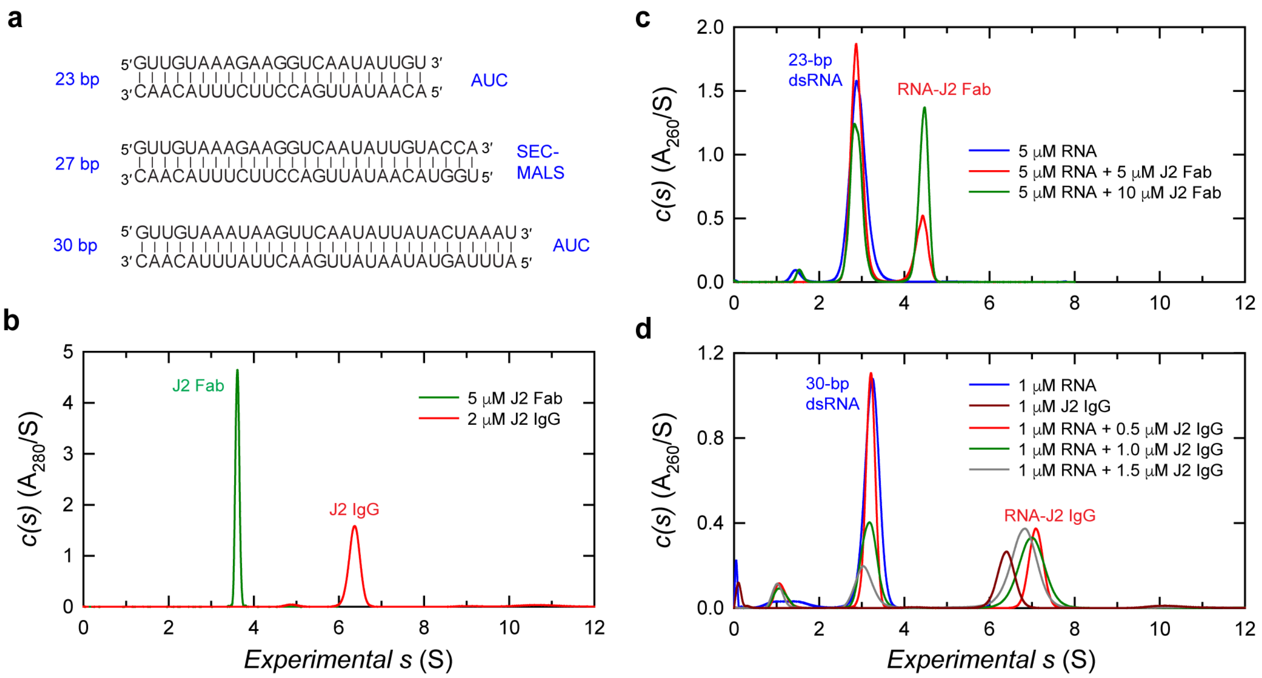

**Supplementary Figure 1. Sedimentation Velocity – Analytical Ultracentrifugation (SV-AUC) analyses of dsRNA interactions with J2 Fab and J2 IgG.** **a**. Sequences of dsRNAs used for AUC and SEC-MALS analyses. **b**. Absorbance c(s) distributions of 5 μM J2 Fab (green) and 2 μM J2 IgG (red). The sample of J2 Fab shows a single species at 3.61 S and 52 kDa indicative of a monomeric Fab. The sample of J2 IgG shows a species at 6.35 S and 147 kDa, indicative of a monomeric IgG. The IgG monomer accounts for 81% of the absorbing sedimenting signal; the dimer (~ 7%) and faster sedimenting material contribute to the signal. **c**. Titration of a 23-bp dsRNA (**a**) with J2 Fab (**b**). The 23-bp dsRNA shows a species at 2.90 S and 14 kDa, indicative of the expected dsRNA duplex. Adding of 1.0 (red) or 2.0 (green) molar equivalents of J2 Fab results in the contribution of a species at 4.39 S and 52 kDa. This species sediments faster than the J2 Fab (3.61 S), indicating a 1:1 RNA:Fab complex. **d**. Titration of a 30-bp dsRNA (17% GC, **a**) with J2 IgG (**b**). The 30-bp dsRNA shows a species at 3.23 S and 19 kDa, indicative of the expected dsRNA duplex. Adding 0.5 equivalents of J2 IgG (red) results in the contribution of a species at 7.09 S and 142 kDa. The species sediments faster than J2 IgG (6.35 S), indicating a 1:1 RNA:IgG complex. The addition of 1.0 (green) or 1.5 (gray) molar equivalents of J2 IgG shows a similar faster sedimenting species with increasing contributions from unresolved free IgG. The plot in brown shows a portion of the c(s) sedimentation profile for 1 μM free J2 IgG. Sample losses were observed due to the formation of large aggregates between the dsRNA and the IgG multimer and faster sedimenting material. Data for (**b**) were collected at 50,000 rpm and 280 nm, whereas data for (**c**) and (**d**) were collected at 50,000 rpm and 260 nm. 12 mm pathlength cells were used in (**b**) and (**d**), whereas 3 mm pathlength cells were used in (**c**).

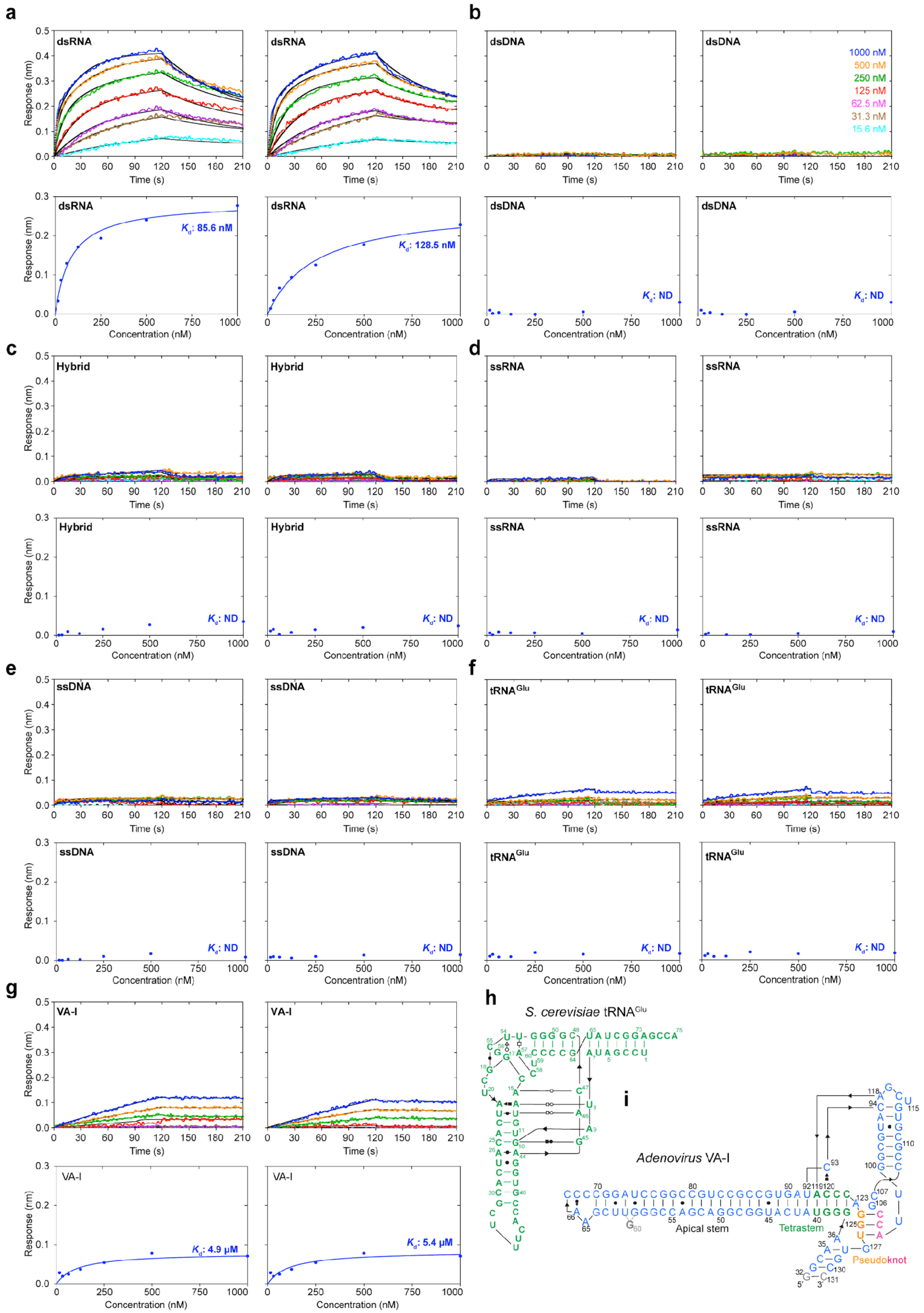

a
b
c
d
e
f
g
h
i
dsRNA
dsDNA
Hybrid
ssRNA
ssDNA
tRNA^Glu
VA-I
Response (nm)
Time (s)
Concentration (nM)
1000 nM
500 nM
250 nM
125 nM
62.5 nM
31.3 nM
15.6 nM
K_d: 85.6 nM
K_d: 128.5 nM
K_d: ND
K_d: 4.9 μM
K_d: 5.4 μM
S. cerevisiae tRNA^Glu
Adenovirus VA-I
Apical stem
Tetrastem
Pseudoknot

**Supplementary Figure 2. J2 IgG binding to different types of nucleic acids. a**-**g**, Additional BLI sensorgrams of J2 IgG binding to dsRNA (**a**), dsDNA (**b**), RNA-DNA hybrid (**c**), ssRNA (**d**), ssDNA (**e**), $tRNA^{Glu}$ (**f**), or VA-I (**g**) at 400 mM KCl. **h**. Secondary structure of $tRNA^{Glu}$ used in **f**. **i**. Secondary structure of VA-I used in **g**. Other nucleic acids used in (**a**-**f**) are shown in Fig. 1a.

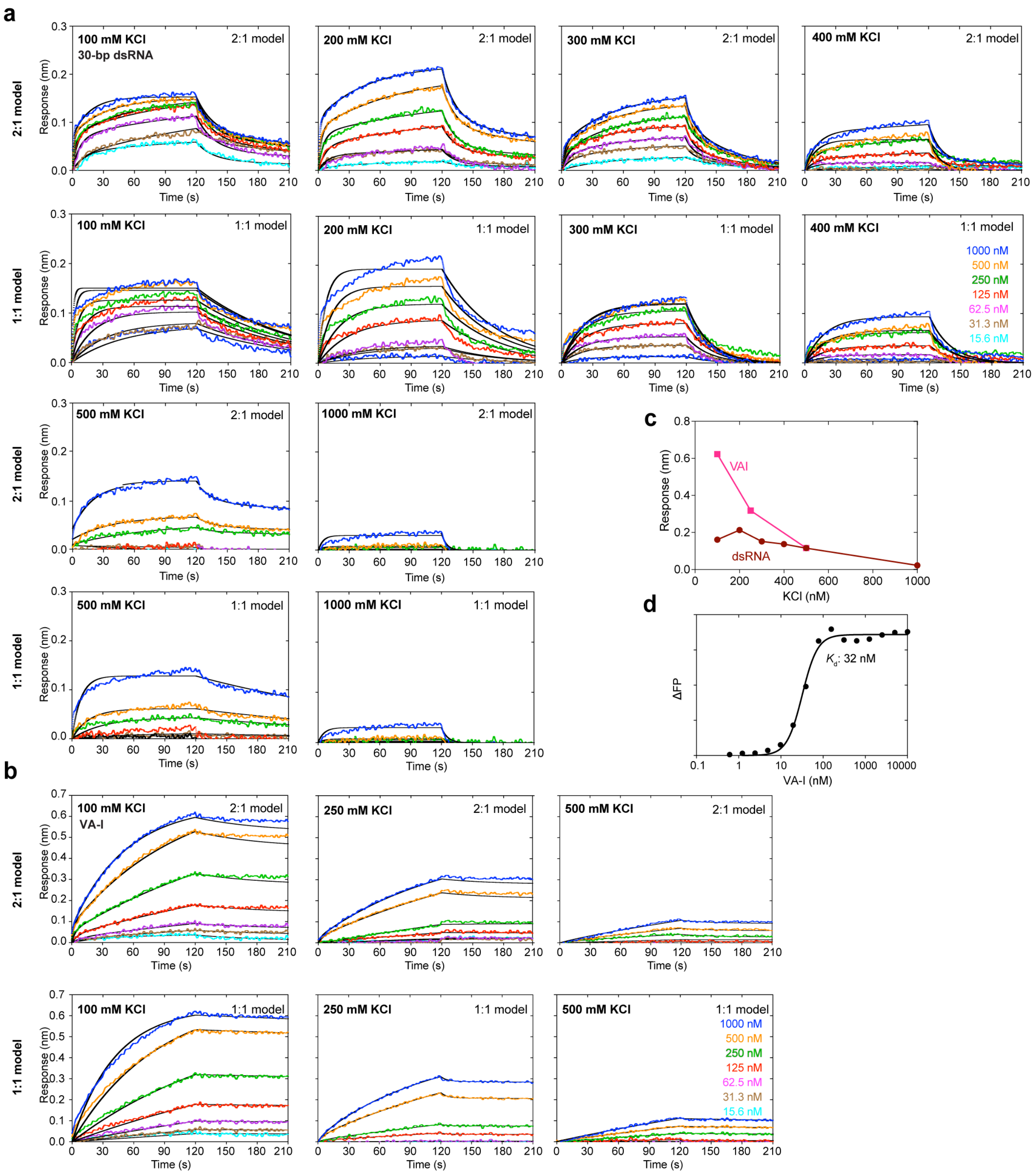

**Supplementary Figure 3. Effects of ionic strengths on J2 IgG binding to dsRNA or VA-I. a**, BLI sensorgrams of J2 binding to a 30-bp dsRNA, at 100, 200, 300, 400, 500, or 1000 mM KCl, fit to the 2:1 model (heterogeneous ligand, upper), or 1:1 model (lower). **b**. BLI sensorgrams of J2 binding to VA-I RNA at 100, 250, or 500 mM KCl, fit to the 2:1 model (upper), or 1:1 model (lower). **c**, Plots of maximum sensor responses as functions of KCl concentrations, derived from **a** & **b**. **d**, Additional bimolecular fluorescence polarization (FP) analysis of J2 IgG binding to a 3′-FAM-labelled VA-I RNA. ΔFP: change in FP expressed in dimensionless millipolarization (mP) units.

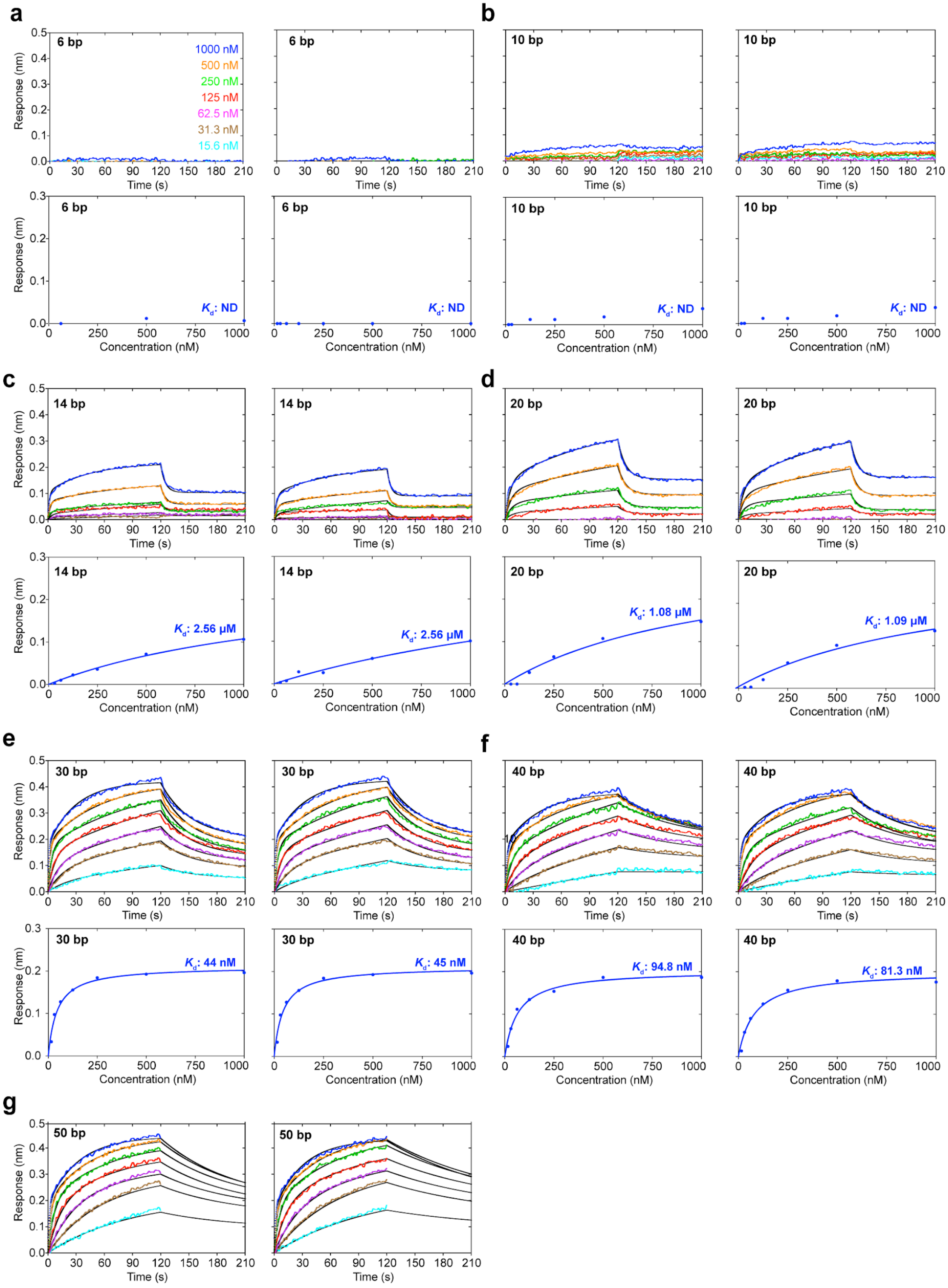
a
b
c
d
e
f
g
6 bp
10 bp
14 bp
20 bp
30 bp
40 bp
50 bp
1000 nM
500 nM
250 nM
125 nM
62.5 nM
31.3 nM
15.6 nM
Response (nm)
Time (s)
Concentration (nM)
$K_d$: ND
$K_d$: 2.56 μM
$K_d$: 1.08 μM
$K_d$: 1.09 μM
$K_d$: 44 nM
$K_d$: 45 nM
$K_d$: 94.8 nM
$K_d$: 81.3 nM

**Supplementary Figure 4. J2 IgG binding to dsRNAs of different lengths. a-g**, Additional BLI sensorgrams (upper) and steady-state analyses (lower) of J2 IgG binding to 6 (**a**), 10 (**b**), 14 (**c**), 20 (**d**), 30 (**e**), 40 (**f**), or 50 bp (**g**) dsRNAs. Sequences shown in Fig. 1e. Due to significant dependency of binding on ionic strength, 100 mM KCl was used for 6, 10, 14, and 20 bp dsRNAs; 200 mM KCl for 30 bp dsRNA; 400 mM KCl for 40 and 50 bp dsRNAs.

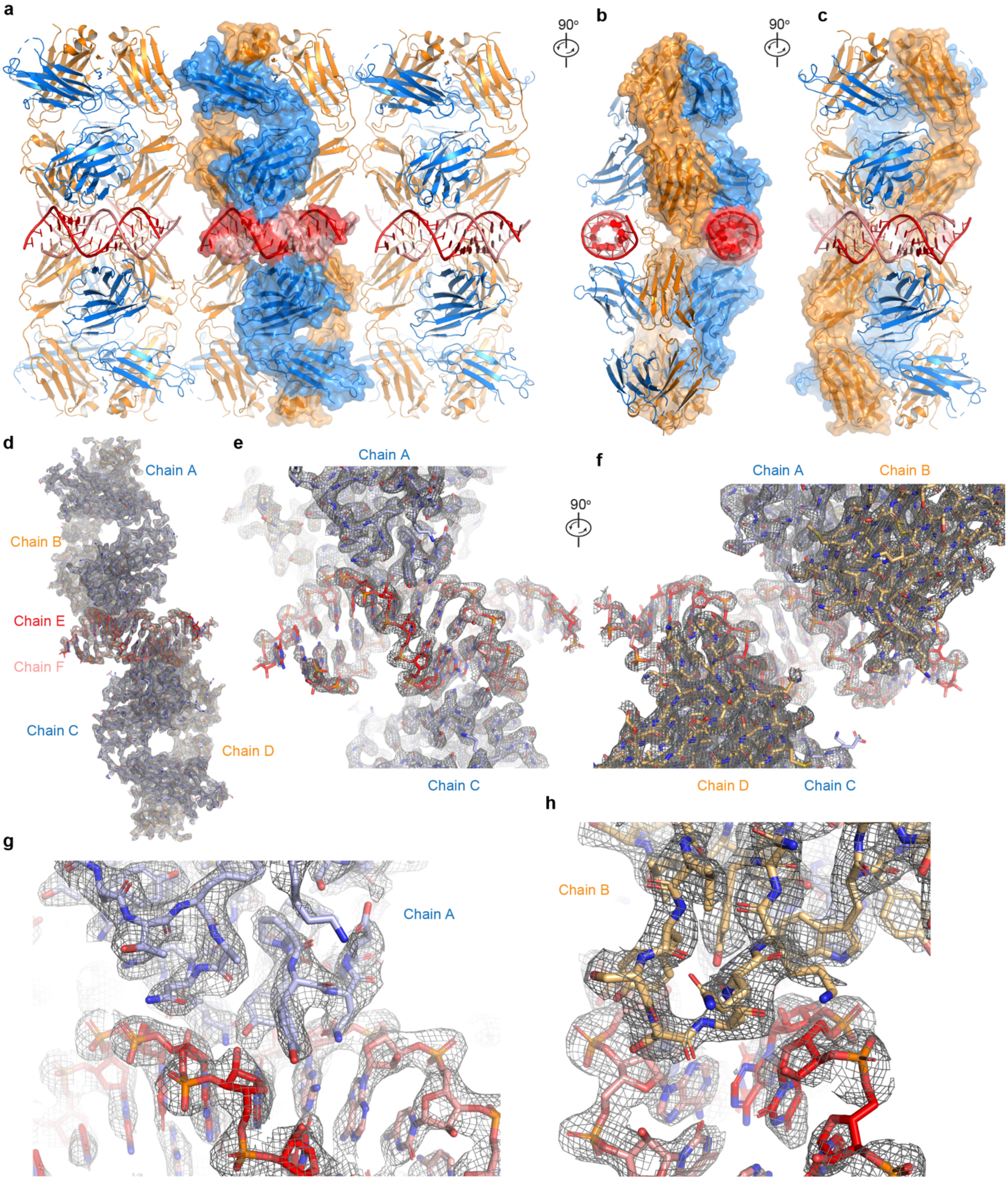

a
90°
b
90°
c
d
Chain A
Chain B
Chain E
Chain F
Chain C
Chain D
e
Chain A
Chain C
f
90°
Chain A
Chain B
Chain D
Chain C
g
Chain A
h
Chain B

**Supplementary Figure 5. Crystal packing arrangements and representative electron densities.** **a**, Crystal-packing arrangements of the J2 Fab – dsRNA (2:1) molecular assemblies. The heavy and light chains of the J2 Fabs are shown in blue and orange, respectively. The two dsRNA strands are in red and salmon. Contents within a single asymmetric unit are highlighted in translucent surface representation. **b**, 90° rotated view of **a**, showing two asymmetric units. **c**, 90° rotated view of **b**, showing two asymmetric units, and a rear view relative to **a**. **d**, Composite simulated anneal-omit 2Fo-Fc density calculated using the final model of two J2 Fabs bound to a 23-bp dsRNA, superimposed with the final refined model, contoured at 0.8 $\sigma$. **e**, **f**, Two views of a portion of the map in **d** showing the J2 interface with the dsRNA strands. **g**, Portion of the map in **d** showing the J2 heavy chain interface with the dsRNA. **h**, Portion of the map in **d** showing the J2 light chain interface with the dsRNA.

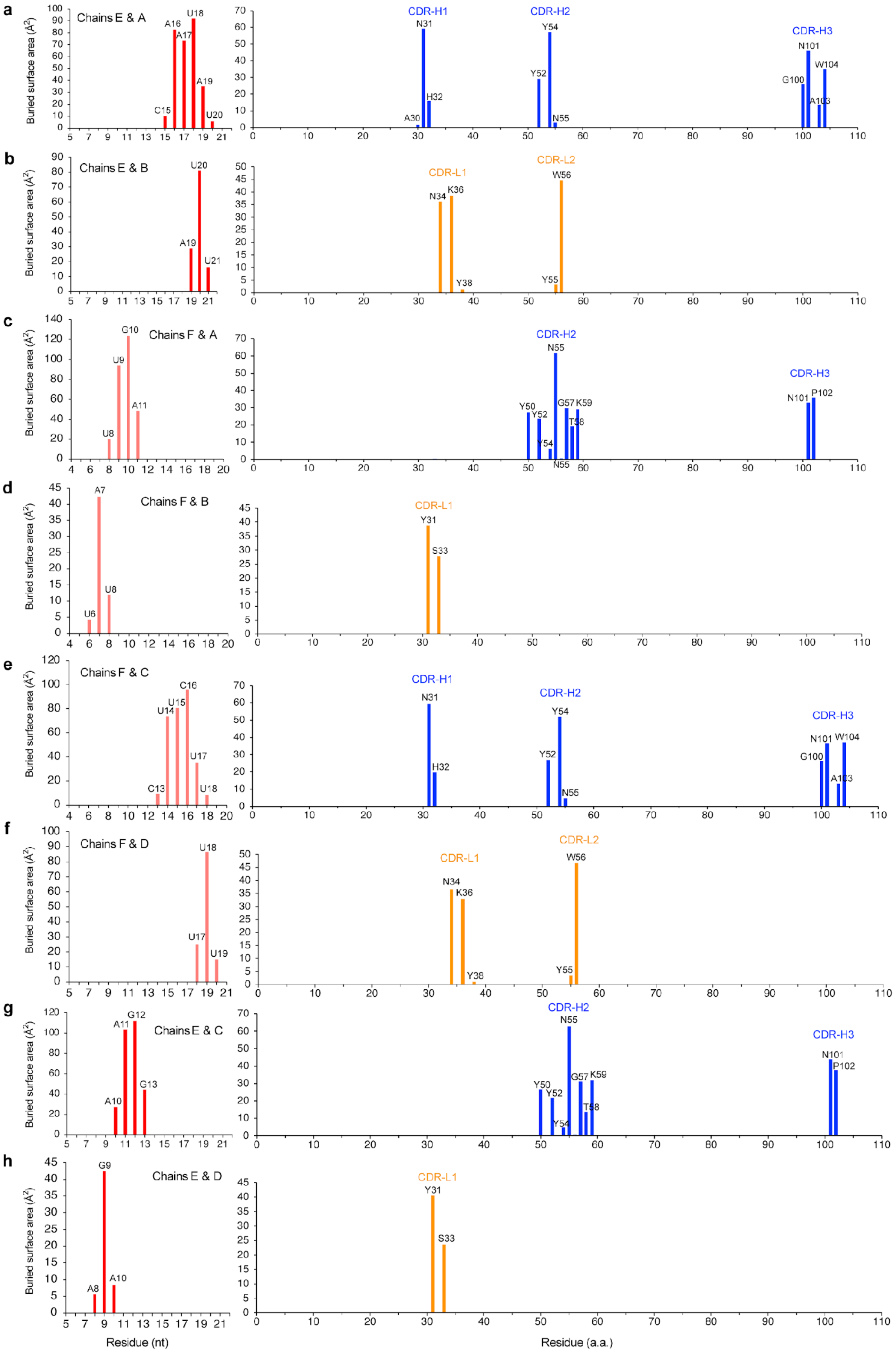
a
Chains E & A
C15
A16
A17
U18
A19
U20
CDR-H1
A30
N31
H32
CDR-H2
Y52
Y54
N55
CDR-H3
G100
N101
A103
W104
b
Chains E & B
A19
U20
U21
CDR-L1
N34
K36
Y38
CDR-L2
Y55
W56
c
Chains F & A
U8
U9
G10
A11
CDR-H2
Y50
Y52
Y54
N55
G57
T58
K59
CDR-H3
N101
P102
d
Chains F & B
U6
A7
U8
CDR-L1
Y31
S33
e
Chains F & C
C13
U14
U15
C16
U17
U18
CDR-H1
N31
H32
CDR-H2
Y52
Y54
N55
CDR-H3
G100
N101
A103
W104
f
Chains F & D
U17
U18
U19
CDR-L1
N34
K36
Y38
CDR-L2
Y55
W56
g
Chains E & C
A10
A11
G12
G13
CDR-H2
Y50
Y52
Y54
N55
G57
T58
K59
CDR-H3
N101
P102
h
Chains E & D
A8
G9
A10
CDR-L1
Y31
S33
Buried surface area (Å$^2$)
Residue (nt)
Residue (a.a.)

**Supplementary Figure 6. Solvent-accessible surfaces buried at the J2 Fab – dsRNA interfaces.** **a**-**d**, Plots of solvent-accessible surface area (Å$^2$) buried per residue on the dsRNA strands (left, chain E or F) and the first J2 Fab (right, heavy chain A in blue; light chain B in orange). **e**-**h**, Plots of solvent-accessible surface area (Å$^2$) buried per residue on the dsRNA strands (left, chain F or E) and the second J2 Fab (right, heavy chain C in blue; light chain D in orange).

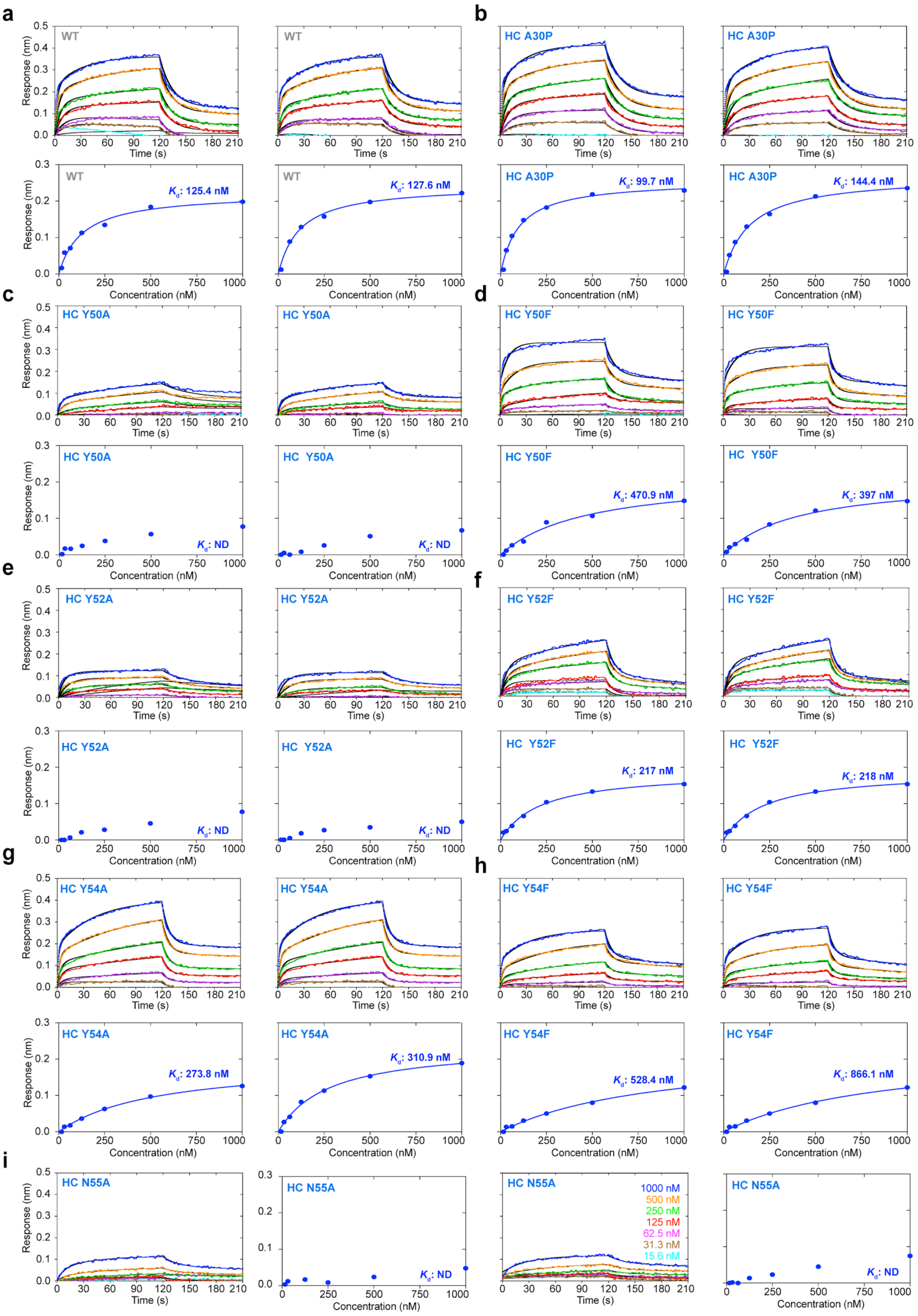
a
WT
WT
Response (nm)
Time (s)
Concentration (nM)
Kd: 125.4 nM
Kd: 127.6 nM
b
HC A30P
HC A30P
Kd: 99.7 nM
Kd: 144.4 nM
c
HC Y50A
HC Y50A
Kd: ND
Kd: ND
d
HC Y50F
HC Y50F
Kd: 470.9 nM
Kd: 397 nM
e
HC Y52A
HC Y52A
Kd: ND
Kd: ND
f
HC Y52F
HC Y52F
Kd: 217 nM
Kd: 218 nM
g
HC Y54A
HC Y54A
Kd: 273.8 nM
Kd: 310.9 nM
h
HC Y54F
HC Y54F
Kd: 528.4 nM
Kd: 866.1 nM
i
HC N55A
HC N55A
Kd: ND
HC N55A
HC N55A
Kd: ND
1000 nM
500 nM
250 nM
125 nM
62.5 nM
31.3 nM
15.6 nM

**Supplementary Figure 7. Mutational analyses of the J2-dsRNA interface, part 1. a-i**, Additional BLI sensorgrams (upper or left) and steady-state analyses (lower or right) of WT and mutant J2 IgG binding to a 20-bp dsRNA shown in Fig. 1e, at 100 mm KCl

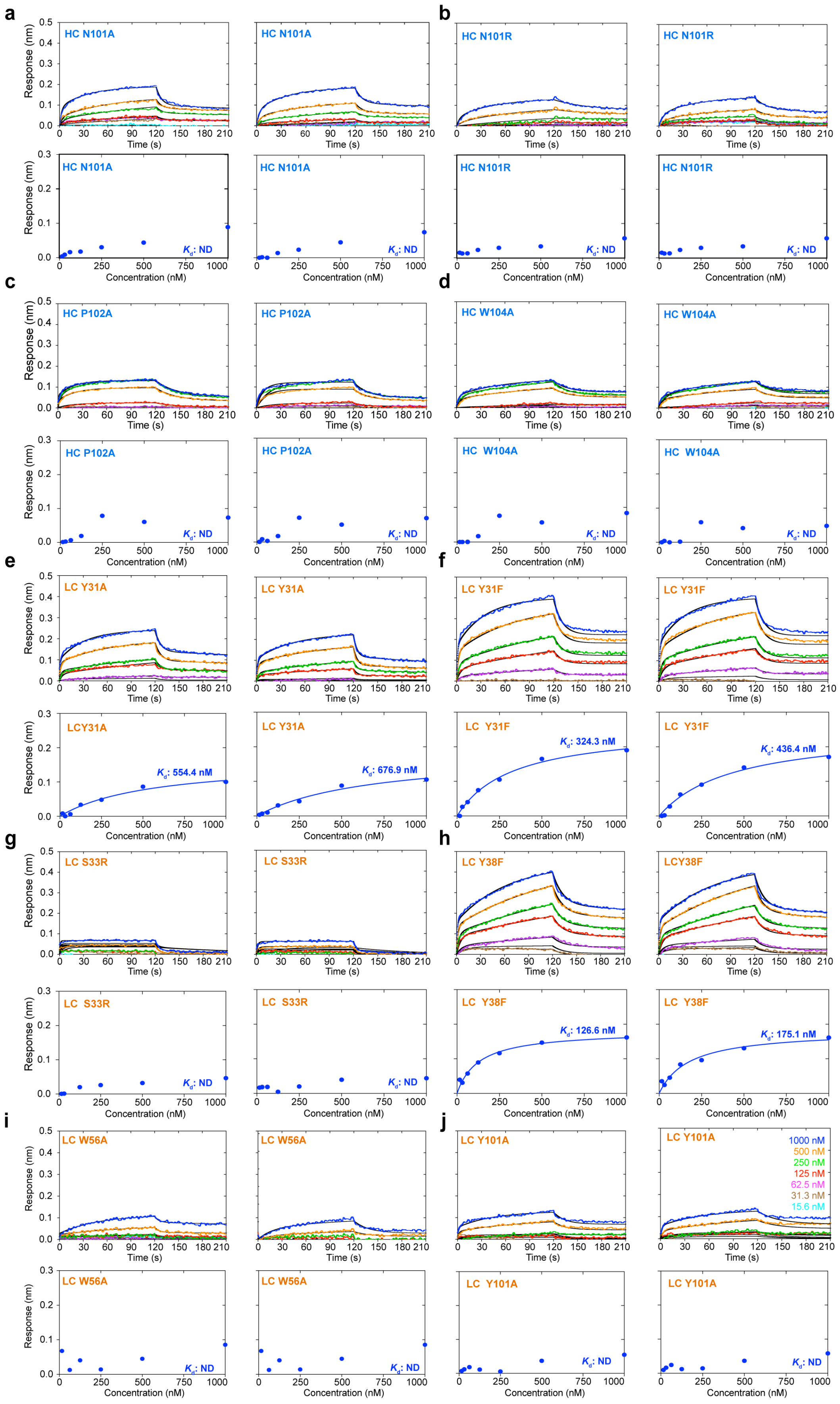

a
HC N101A
b
HC N101R
c
HC P102A
d
HC W104A
e
LC Y31A
LCY31A
f
LC Y31F
g
LC S33R
h
LC Y38F
LCY38F
i
LC W56A
j
LC Y101A
Response (nm)
Time (s)
Concentration (nM)
Kd: ND
Kd: 554.4 nM
Kd: 676.9 nM
Kd: 324.3 nM
Kd: 436.4 nM
Kd: 126.6 nM
Kd: 175.1 nM
1000 nM
500 nM
250 nM
125 nM
62.5 nM
31.3 nM
15.6 nM

**Supplementary Figure 8. Mutational analyses of the J2-dsRNA interface, part 2. a-j**, Additional BLI sensorgrams (upper) and steady-state analyses (lower) of WT and mutant J2 IgG binding to a 20-bp dsRNA shown in Fig. 1e, at 100 mm KCl.

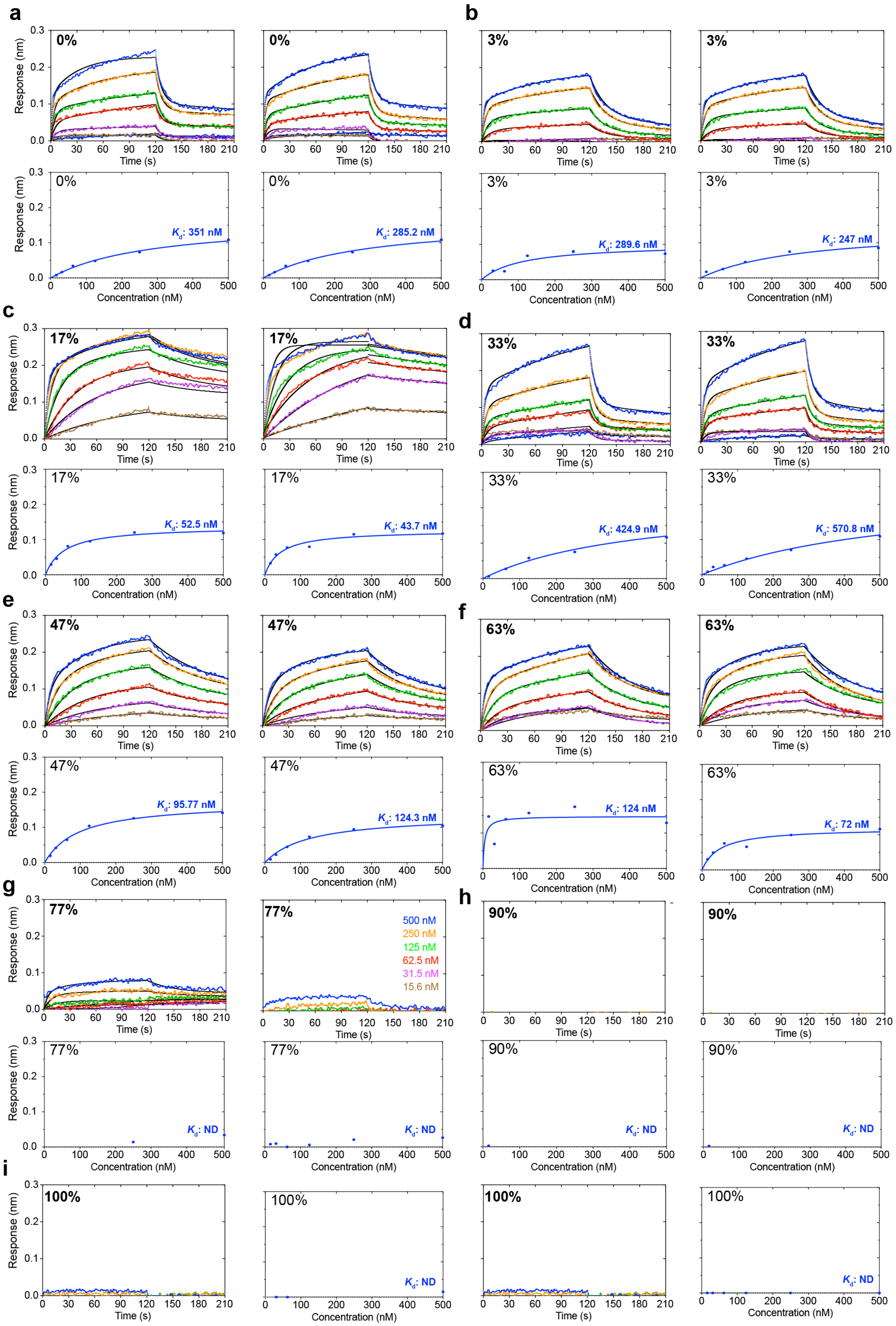
a
b
c
d
e
f
g
h
i
0%
3%
17%
33%
47%
63%
77%
90%
100%
Response (nm)
Time (s)
Concentration (nM)
Kd: 351 nM
Kd: 285.2 nM
Kd: 289.6 nM
Kd: 247 nM
Kd: 52.5 nM
Kd: 43.7 nM
Kd: 424.9 nM
Kd: 570.8 nM
Kd: 95.77 nM
Kd: 124.3 nM
Kd: 124 nM
Kd: 72 nM
Kd: ND
500 nM
250 nM
125 nM
62.5 nM
31.5 nM
15.6 nM

**Supplementary Figure 9. Effects of GC contents on J2 IgG binding. a-i**, Additional BLI sensorgrams (upper or left) and steady-state analyses (lower or right) of J2 binding to 30-bp dsRNAs shown in Fig. 4a, with GC contents of 0% (**a**), 3% (**b**), 17% (**c**), 33% (**d**), 47% (**e**), 63% (**f**), 77% (**g**), 90% (**h**), or 100% bp (**i**), at 200 mM KCl.

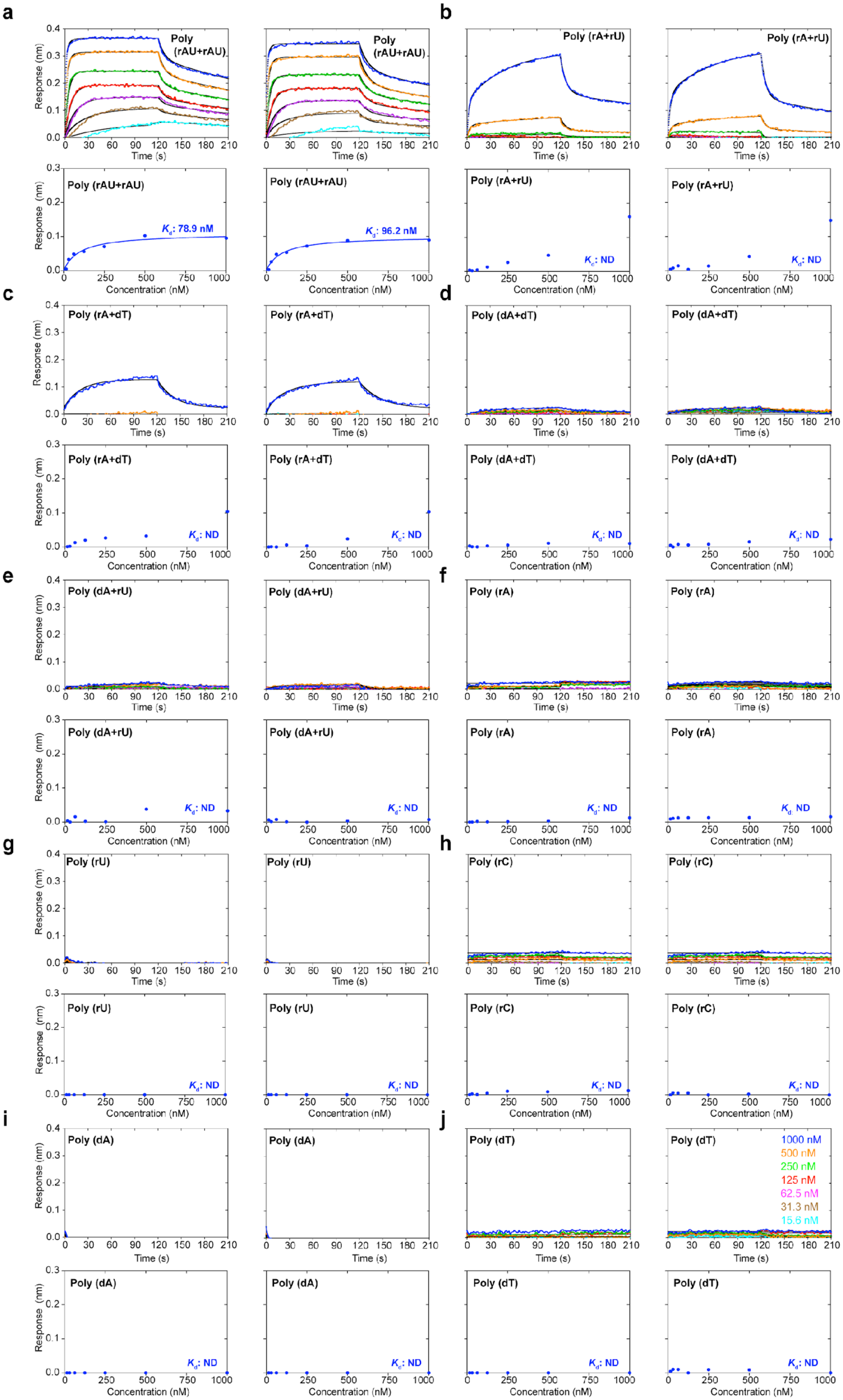
a
b
c
d
e
f
g
h
i
j
Poly (rAU+rAU)
Poly (rA+rU)
Poly (rA+dT)
Poly (dA+dT)
Poly (dA+rU)
Poly (rA)
Poly (rU)
Poly (rC)
Poly (dA)
Poly (dT)
$K_d$: 78.9 nM
$K_d$: 96.2 nM
$K_d$: ND
Response (nm)
Time (s)
Concentration (nM)
1000 nM
500 nM
250 nM
125 nM
62.5 nM
31.3 nM
15.6 nM

**Supplementary Figure 10. J2 IgG interaction with homotypic repeat nucleic acid sequences.** **a-j**, Additional BLI sensorgrams (upper) and steady-state analyses (lower) of J2 binding to repetitive nucleic acid sequences shown in Fig. 5a, at 400 mM KCl.
